# Assessing Long-Read Mappers for Viral Genomics

**DOI:** 10.1101/2024.11.25.625163

**Authors:** Thomas Baudeau, Camille Marchet, Mikaël Salson

## Abstract

Long-read sequencing technologies from Pacific Biosciences and Oxford Nanopore Technologies (ONT) have advanced genomic research, producing reads over 10 kilobases and enabling rapid field-based viral surveillance. This study evaluates eight long-read mapping tools on viral genomic data, including modern and legacy methods. We assessed their performance on ONT reads and their impact on variant calling using bcftools and medaka. Using simulated and real datasets under varying conditions, reflecting different experimental and biological conditions, such as variable read lengths, error rates and the presence of multiple viral variants, we found that the majority of tools had great difficulty in correctly managing read edges. In addition, it was found that with a default setting, the performance of the tools decreased.

## INTRODUCTION

The introduction of long-read sequencing technologies, such as those developed by Pacific Biosciences ^1^ (PacBio) and Oxford Nanopore Technologies ^2^ (ONT), has significantly improved our capacities in comparison to short-read sequencing. These long-read technologies can produce sequences exceeding 10 kilobases in length on average, substantially enhancing the accuracy and confidence in genome assembly ^3^. Notably, ONT can generate reads up to 1,000 kb in length with an accuracy varying from 85 % to close to 100 % ^4^. Despite early limitations such as lower throughput, higher error rates, and greater cost per base, recent advancements have improved these aspects considerably ^5^.

ONT’s portable sequencing devices facilitate rapid, on-site genome sequencing from environmental or clinical samples, eliminating the need for extensive laboratory infrastructure ^6^. This capability has been a game changer in monitoring viral outbreaks, as exemplified by the ARTIC protocols ^7^, which leverages ONT reads for field-based viral surveillance. Similar to the ARTIC approach, studies utilizing long-read sequencing for viral research use mapping techniques to align reads to known genomes and identify variants or other genomic features. Long-read mappers ^8–12^, which are tailored to handle the specific error profiles and lengths of these reads, have predominantly been developed for well-characterized eukaryotic genomes. However, they are often applied to other organisms using default parameters ^13^. Although benchmarks for mapping quality typically use eukaryotic and prokaryotic genomes ^14,15^, they rarely include viral species.

Viral genomes, which are compact enough to be covered by a single long read in some cases, present however unique challenges. Actually, many viral sequencing protocols, such as the aforementioned ARTIC ones, rely on tiled amplicons in order to cover the whole genome, resulting in read lengths much shorter (around 500 bases) than those used in most eukaryotic applications. Additionally, the high mutation rates and diversity within viral populations can complicate mapping efforts ^16^. Viruses also produce a diversity of sequences varying from species to species and linked to singular molecular mechanisms, such as *splicing* in HIV ^17^ or *subgenomic RNAs* for SARS-CoV ^18^. These sequences can be different enough from the reference to be a challenge for mappers to locate them accurately with default settings ^19^. Nevertheless, the much smaller size of viral genomes compared to eukaryotic genomes should allow for more computational resources to be allocated to achieve higher accuracy. Interestingly, while modern mappers are designed for speed, older methodologies were slower but provided a more thorough use of the available sequence data. This discrepancy raises the question of whether earlier mapping techniques might be better suited for the unique characteristics of viral genomes. Currently, there is a lack of comprehensive studies comparing mapping results across various viral genomes.

## MATERIALS AND METHODS

### Choice of mapping tools

In this study, we focused on 8 mapping tools and their ability to align ONT reads from viruses. The selection of tools was made in an attempt to diversify the approaches used for mapping and alignment as much as possible. The decision to exclusively focus on ONT data was due to the fact that ONT, notably with the MinION, offers portability well suited during an epidemic ^7^. An *a priori* categorization would lead to separate older short-read approaches (often called seed-and-extend due to their methodology), early long-read approaches (without sampling), and recent long-read approaches (with sampling, and with a seed-chain-extend paradigm). Seeding refers, to the collection of subsequences called “seeds” within a sequence. In the case of short-read mappers, these seeds are directly “extended” through alignments to the genome. For long-read mappers, another step is involved: chaining, which aims at selecting high-scoring chains of small sequence matches between the seed and the reference. This step was computationally relevant to enhance and speed up the base-level alignment (the extension part). In addition, with the emergence of different types of seeds, sampling methods have been developed to collect only a portion of the seeds. Another aspect of mapping, known as soft clipping, will also be examined. Soft clipping consists in not aligning the ends of the reads. All mappers use soft clipping to varying degrees. It helps manage, for example, chimeric reads or the presence of barcodes that may not have been trimmed.

For details on the different algorithms, we report the interested readers to our recent methodological review ^12^.

For the short-read mapping approaches, we selected **bwa-mem2** ^20^ and **magic-BLAST** ^21^. bwa-mem2 was originally designed for short-read data but now incorporates options for long-read mapping. Magic-BLAST is based on BLAST ^22^, using exact fixed-length seeds, but accommodating for indels typical of ONT sequencing errors, and for large gaps generated for instance, but not only, by splicing.

For the early long-read approaches, we included **BLASR** ^23^ and **GraphMap** ^24^. BLASR is one of the first mappers designed for aligning long reads. BLASR indexes all subsequences of size *k* (k-mers) in order to find primary matches between the sequence and the reference (the seeding part). BLASR introduced the “chain” step in the seed-chain-extend paradigm. As BLASR, GraphMap is no longer maintained but is an interesting approach as it indexes all k-mers of the reference while allowing a certain level of substitutions, insertions and deletions, that can increase its sensitivity during the seeding step.

For the recent long-read approaches, we selected **minimap2** ^8^, **Winnowmap2** ^10^, **BLEND** ^25^ and **lra** ^11^. minimap2 is a popular choice for various bioinformatics applications, including genome assembly, variant calling, and metagenomics analysis. minimap2 is integrated in several pipelines or tools like periscope ^13^ or the ARTIC pipeline. It also integrates various novelties presented in different papers. Like the other recent approaches, it harvests putative matches using a sampling approach based on minimizers. Winnowmap2 is a long-read mapper based on Winnowmap2^9^. BLEND is a long-read mapper specialized in high quality long-read data. lra is a long-read mapper that has proposed a different chaining algorithm to better manage the alignment of structural variants, and obtained the best results in some datasets of a recently published benchmarks ^26^.In addition to the previously mentioned tools, an older version of minimap2 was added. Our goal was to analyze and better understand how successive versions of a tool impact its performance. In addition to this version, a version integrating a particular type of seed has been added. This version will be called syncmer-minimap2^27^.

#### Impact of mapping tools on downstream analyses

In order to assess the impact of mappers’ parameters on the rest of the analysis, we chose to focus on a classic application: variant calling. We selected two variant callers, bcftools ^28^ and medaka (github.com/nanoporetech/medaka). We chose medaka as it is included in the ARTIC pipeline, for viral ONT data ^7^. Additionally, medaka is specialized in haploid variant detection, rendering it the most suitable tool for our investigation. medaka variant detection relies on neural networks. We used the two models available on the medaka GitHub repository, R9.4 and R10.3, that reflect the ONT pore versions. Upon scrutinizing the available variant callers, we noted that many now incorporate components involving pre-trained or trainable neural networks. Given the challenges in acquiring viral data and the ensuing difficulty in training, the prevalence of pretrained models Given the difficulties of acquiring viral data and the consequent difficulties of training, the prevalence of pre-trained models means that we have easy access to a tool whose effectiveness on the species on which it has been trained is well known. However, the opaque nature of the training methodologies for such models raises concerns regarding sensitivity or bias. Therefore, the inclusion of a variant caller that eschews such methods is deemed essential to identify potential training biases in certain mappers among the variant callers, hence the inclusion of bcftools.

#### Comparing and filtering the vcf

In order to compare the results provided by the two variant callers, two methods have been implemented. The first method uses vcfdist ^29^ which has already been used to compare variants calling tools ^30^ for bacteria. In addition to this tool, a much simpler home-made comparison method was created. This method checks, for each variant found, whether an identical variant exists within a margin of ±1 nucleotide. Moreover, unlike vcfdist, this method ignores the quality of the variant detection (as provided by the QUAL tag in the VCF files). Additionally, for both tools, the VCF files will be filtered, and all variants located within the first 30 nucleotides, as well as any variants occurring within 10 nucleotides before the poly-A tail, will be excluded. This was added because vcfdist encountered issues when handling variants at the edges of the reads.

#### Generation of ground-truth datasets

Accurate mapping and variant calling assessment requires to use various datasets for which the ground truth is known. We generated many samples and their corresponding sequencing datasets *in silico*. We wanted to mimick different experimental and biological settings. We mainly anticipated the following difficulties in the case of viral long-read data. Some difficulties come from preparation conditions.The reads size and coverage can vary according to the protocol for the size, it’s influenced by the sequencing protocol (*eg*. amplicon or shotgun), and by wet-lab conditions. The coverage depends on factors such as the quantity of material to initiate the sequencing, the level of contamination and the capacity to amplify virus-specific sequences. Biologic modalities also come into place: viruses have the faculty to mutate a lot in a given host, leading to the presence of variants and quasispecies in the samples. Depending on the aim of the experiment, specific viral sequences can also appear, such as subgenomic RNAs which combine separate parts of the viral genome.

In order to have results that encompass the different data specificities according to the protocols. We generated samples from 10 different experimental conditions. The ONT datasets were simulated using pbsim2^31^, using variable parameters to set the simulated sequencing depth (`--depth`), the distribution of the read lengths (`--length-mean, --length-min, --length-max`), the sequencing accuracy (`--accuracy-mean`) and the HMM model corresponding to the sequencer being simulated (`--hmm_model`). We also set a fixed parameter for the ratio or substitutions, insertions and deletions, as recommended by pbsim2 documentation for Nanopore (`--difference-ratio 23:31:46`). Each dataset has a variety of conditions, reflecting the diversity of technical or biological aspects. The conditions are as follows:

- **Length**: the average size of reads in the dataset.
- **Virus**: the reference virus in the dataset
- **Error Rate**: the error rate in each dataset
- **Model**: the pore model to be copied by pbsim2
- **Contamination**: the percentage of human reads that will be added to the sample
- **Coverage**: the coverage for each dataset
- **Nb of variants**: the number of variants present in the sample
- **Proportion of variants**: the percentage of reads originating from this variant
- **Variations**: the number of mutations introduced for each variant

We define eight datasets, each varying by some of the aforementioned conditions in order to assess their effects on the mapping or on the variant calling. All parameters and names for each dataset are presented in table 1. We summarize below the characteristics we wanted to asses in each dataset:

**Table 1:** Table of the different conditions for each experiment.

| Name | Length | Species | Error Rate | Model | Contamination (%) | Coverage | nb Variant | prop Variant | %v Variant |
| --- | --- | --- | --- | --- | --- | --- | --- | --- | --- |
| D1 | 500,7500 | covid | 0.95 0.99 0.90 | R103 | 10 | 10 | 1 | 99 | 2 |
| D2 | 500,7500 | covid,vih | 0.95,0.99,0.90 | R103 | 10 | 100 | 1 | 99 | 2 |
| D3 | 500,7500 | covid | 0.95 0.99 0.90 | R103 | 10 | 1000 | 1 | 99 | 2 |
| D4 | 500,7500 | covid | 0.95 | R103 | 10000 | 100 | 1 | 99 | 2 |
| D5 | 500,7500 | covid | 0.95 | R103 | 10 | 100 | 1 | 99 | 7 |
| D6 | 500,7500 | covid | 0.95 | R94 | 10 | 100 | 1 | 99 | 2 |
| D7 | 500,7500 | covid | 0.95 | R94/R103 | 10 | 100 | 1 | 99 | 2 |
| D8 | 500,7500 | covid | 0.95 | R94 | 10 | 100 | 1 | 50 | 2 |

**D1** low sequencing coverage

**D2** different viruses

**D3** high sequencing coverage

**D4** high host contamination of the sample

**D5** high divergence between the sequenced virus and the reference one

**D6**,**D7** different sequencing pores

**D8** two similar viruses in the sample

### Real datasets

In addition to the generated datasets, 4 real datasets are used. The 4 datasets, taken from a previous study ^32^, can be found at the following address: github.com/iqbal-lab-org/covid-truth-datasets or on the ENA project with the IDs PRJEB51850 and PRJEB50520. In this study, the SARS-CoV-2 variants were detected with high quality sequencing runs from both Illumina and ONT sequencing, each variant has been manually validated. We take only 4 datasets from the study, we choose variant dataset where ARTIC sequencing run was available. The 4 datasets will be named R1a, R1b, R2a, and R2b, and they were all sequenced using a Nanopore platform. R1a consists of 55,615 reads sequenced following the midnight-v2 amplicon scheme. R1b consists of 557,718 reads following the artic-v4.1 amplicon scheme. Both datasets were sequenced from the same strain, which has 4 substitutions and one deletion. R2a consists of 45,484 reads sequenced following the midnight-v2 amplicon scheme. R2b consists of 157,189 reads following the artic-v4.1 amplicon scheme. Both datasets were sequenced from the same strain, which has 32 substitutions and 3 indels. All datasets contain barcodes.

### Pipeline of the benchmark

We developed a fully-reproducible pipeline in Python (https://github.com/ThomasBaudeau/BenchMapL) to generate ONT data and compare all the mapping tools previously introduced as well as the variant callers. The pipeline is built using Snakemake ^33^ and consists of several steps. The first step consists in creating a reference sequence for each variant. This step uses a reference FASTA input file and produces a FASTA file for each variant as well as a VCF file listing all the mutations introduced in the variants. The mutations can either be short indels (one or two bases) or SNPs. The second step, consists in generating the high-throughput sequencing dataset for each variant with pbsim2^31^. It produces a FASTQ file for each variant and the expected alignment on the reference of the corresponding variant. During the third step, those expected alignments are then corrected to represent the alignment on the original reference. These corrected alignments will be referred to as “perfect” or “perfect BAM” in the rest of the paper. In the fourth step, all the FASTQ files are merged together depending on the proportions for each variant provided by the user. Additional contamination from human reads can also be added, the reads come from sample ID SRR16071311 available on NCBI SRA. The next step is to run all the mappers. Then, the output files are processed using SAMTOOLS and plots on some metrics are generated. During the last step, the variant callers are launched and generate a VCF file. The resulting files are compared with the expected one. All the metrics are calculated and stored in a CSV output file. The various computed metrics for the comparison are :

- **% of mapped reads (M)** 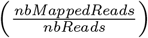: This corresponds to the number of reads from the species mapped. The added human reads are excluded from the percentage.
- **% of soft-clipped bases (sB)** 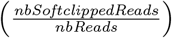: This corresponds to the sum of soft clipped read.
- **% of shifted bases (scB)** 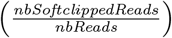: This corresponds to the sum of the offset between the expected positions of the first and last nucleotides and the observed position.This metric is only used for simulated data
- **number of contaminated reads mapped (nC)**: Number of human reads mapped. number of wrongly mapped read this metric is only used for simulated data
- **False positive (FP)**: Number of variants incorrectly called.
- **False negative (FN)**: Number of missed variants.
- **True positive (TP)**: Number of corrects variants found.
- **F1-score**:

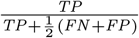

### Parametrizing the mapping tools

For each tool, there are three sets of parameters. In most cases, the parameters that are modified will affect the indexing process. The goal is to determine whether these changes will impact the quality of the mapping or only the execution speed of the tools. All the tool authors were contacted and asked which parameter settings would be most appropriate for each tool. The parameters recommended by the authors are marked with an asterisk in the table 2.

**Table 2:** Table of used parameter settings for each tool.

| tool name | Version | Parameter details | Default parameter | Parameter One | Parameter Two |
| --- | --- | --- | --- | --- | --- |
| minimap2 | 2.28 | -k the size of the minimizers -w the size of the window | k = 15<br>w = 10 | k = 12<br>w = 4 | k = 7<br>w = 5. |
| minimap2 (old) | 2.1.1 | -k the size of the minimizers -w the size of the window | k = 15<br>w = 10 | k = 12<br>w = 4 | k = 7<br>w = 5. |
| syncmer-minimap2 | 2.1.1 | -k the size of the k-mer, -w the size of the window, -downsample the down-sampling factor, -s-mer the size of the sync-mer, -pos the sync-mer parameters | k = 15<br>w = 10<br>downsample = 1<br>s-mer = 5<br>pos1 = 3<br>pos2 = 9 | k = 12<br>w = 4 | k = 7<br>w = 5. |
| Magic-BLAST | 1.5.0 | -word size , the minimum number of consecutive bases that must match exactly. the minimum value for this parameter is 12. | -word size = 18 | word size = 16 | word size =22 |
| GraphMap | 0.5.2 | The parameter k corresponds to the size of k-mers inside the graph, while A corresponds to the minimal size of a match to be chosen as an anchor. | k = 6<br>A = 12 | k = 7<br>A = 8 | k = 5<br>A = 7 |
| BLASR | 5.3.5 | -minMatch, which corresponds to the seed size, and -nCandidates, which affects the number of alignments tested. | minMatch =12<br>nCandidates =10 | minMatch =12<br>nCandidates =10 | minMatch =12<br>nCandidates =10 |
| bwa-mem2 | 0.5.2 | -k minimum seed length; -W discard a chain if seeded bases shorter than W; -w band width for banded alignment | k = 14<br>w = 1<br>W = 1, | k = 7<br>W = 10<br>w = 1 | k = 5<br>w = 1<br>W = 1 |
| Winnowmap2 | 2.03 | -k , which correspond to the size of the k-mer, and -w, which corresponds to the size of the window, are used. | k = 15<br>w = 50 | k = 15<br>w = 1 | k = 12<br>w = 4 |
| BLEND | 1.0.0 | -k, representing the size of the k-mer, -w, representing the window size, -fixed-bits number of bits used for the hash values of seeds, and -neighbors representing the number of kmer to generate a seed. | k = 7<br>w = 10<br>fixed-bits = 32<br>neighbors = 11; | k = 7<br>w = 5<br>fixed-bits = 24<br>neighbors = 8 | k = 12<br>w = 5<br>fixed-bits = 30<br>neighbors = 10 |
| Ira | 1.3.2 | -W The minimizer window size,-K The word size. | -W =10<br>-k = 17 | -W = 4<br>-k = 12 | -W = 5<br>-k = 7 |

## RESULTS

### General mapping results

#### Raw mapping rates

First, we analyze the mapping rate for each experiment. The tables D1 to D8 in supplementary file 1 and figure 1A show that high mapping rates were observed across almost all tools tested. When combining the 8 experiments, 3 groups of tools can be distinguished (fig 1A). The first group comprises the tools with the best results, including bwa-mem2, GraphMap, minimap2, syncmer-minimap2, BLASR, perfect; the second group corresponds to tools with a lower number of mapped reads in some datasets, made up of lra, BLEND and Winnowmap2; and finally a third group with Magic-BLAST, which on the whole has poorer results than the other tools.

**Figure 1:**
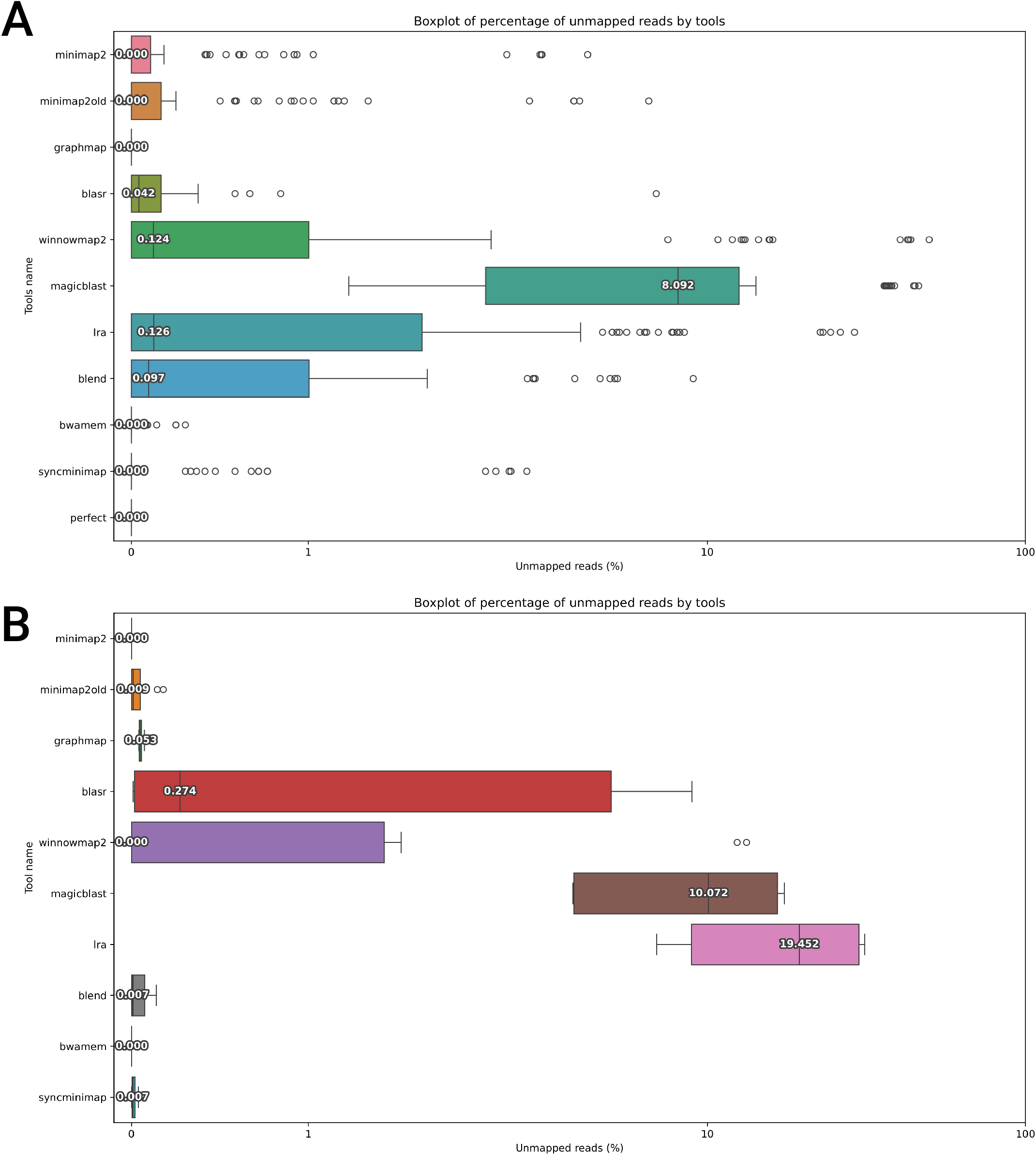
Boxplot of the percentage of 1-M for each tool for the 34 samples generated. The x-axis is in symlog scale.For minimap2, GraphMap,bwa-mem2, syncmer-minimap2, and perfect, the median is 0. For BLASR, it is 0.084, 0.140 for Winnowmap2, 0.195 for BLEND, 0.210 for lra, and 9.575 for Magic-BLAST.

A more detailed analysis of each dataset shows that under conditions replicating ARTIC characteristics (such as read length = 500 and error rate = 10 or 5 %), bwa-mem2 and GraphMap mapped a higher number of reads compared to other approaches under their default settings (Additionnal figures S1 and S2). Furthermore, we also see the impact of error rate and length of the reads on the mapping results. lra, BLEND, Winnowmap2, and Magic-BLAST map less reads when they are shorter, but their results are similar to the other mappers with longer reads. We observe the same effect on error rates, for those tools, with a higher error rate leading to fewer reads mapped.

The results for the average percentage of unmapped reads with real data show similar outcomes for the vast majority of the tools (Figure 1B). However, lra shows a significant decrease in performance, with a median of 19.4% unmapped reads. Magic-BLAST and Winnowmap2 also see an increase in their number of unmapped reads. BLEND, on the other hand, shows improved results with only 0.007% unmapped reads compared to 0.097% with simulated data.

Table 3 shows that on average less than 3% of reads are have their sequence endings and beginnings correctly positioned. We note that two tools, Magic-BLAST and lra, deviate significantly from this average, with 1.34 and 0.07, respectively. We also see that on average more than 10% of the mapped reads have their mapping coordinates shifted by more than 20 nucleotides compared to the theoretical expected mapping.

**Table 3:** Proportion for each tool of the mapped reads according to the shift in mapping coordinates, compared to the theoretical ones, over the 34 experiments. A shift of 0 corresponds to a read that starts and ends at the exact expected positions, a shift of *x* positions means that the start and end have in total a shift of *x* positions compared to the expected theoretical alignment. When measuring the ability of a mapper to correctly map a read, the proportion of reads should be as high as possible for a shift of 0 positions (in green), and the values for the other categories (especially for the last ones) should be as low as possible (the highest values are in red for each categories).

|  | Shift in mapping coordinates |  |  |  |  |
| --- | --- | --- | --- | --- | --- |
|  | 0 | 1-5 | 6-10 | 11-20 | >20 |
| GraphMap | 2.46 | 18.82 | 34.71 | 32.80 | 11.20 |
| BLEND | 2.07 | 17.35 | 33.15 | 34.44 | 12.98 |
| BLASR | 1.77 | 15.36 | 31.25 | 36.87 | 14.74 |
| bwa-mem2 | 1.92 | 15.00 | 28.14 | 27.97 | 26.97 |
| syncmer-minimap2 | 2.07 | 17.41 | 33.22 | 34.47 | 12.83 |
| minimap2 | 2.06 | 17.31 | 33.08 | 34.46 | 13.09 |
| minimap2 (old) | 2.06 | 17.31 | 33.09 | 34.47 | 13.07 |
| Magic-BLAST | 1.34 | 6.50 | 13.01 | 13.74 | 65.40 |
| Winnowmap2 | 2.16 | 17.35 | 33.31 | 34.41 | 12.77 |
| Ira | 0.07 | 3.08 | 9.70 | 30.84 | 56.30 |

In supplementary file S6 and S7 we see that the error rate has no effect of the number of shifted reads, but the length seems to have an effect with a larger number of reads perfectly aligned with a small read size.

Table 4 shows the soft clipping results for each read, indicates that on average, 50% of the reads are not affected by soft clipping. However, certain tools like lra and Magic-BLAST have only 0.93% and 27.53% of their reads, respectively, unaffected by this phenomenon. For GraphMap, 83.49% of the reads show no soft clipping. Figure 2 shows the trends observed for soft clipping on real datasets. We observe on those datasets that, apart from GraphMap and bwa-mem2, the other tools use soft clipping for more than 90% of the reads, with significant soft clipping (greater than 20 bases). It is also noteworthy that bwa-mem2 has 6% of its reads without soft clipping, whereas for the other tools, it’s less than 1%. However, it is important to note that barcode has been present and has not been trimmed on the real data

**Table 4:** The percentage of average soft clipped bases per read for each tools. Each category corresponds to the number of nucleotides soft-clipped in the whole read at either end. The higher the percentage in the column for 0 nucleotide soft-clipped (in green), the better the ability of the mapper to correctly align the read. The values in the other columns should be as low as possible (the highest value is in red for each category).

|  | Nb bases soft-clipped |  |  |  |  |
| --- | --- | --- | --- | --- | --- |
|  | 0 | 1-5 | 6-10 | 11-20 | >20 |
| GraphMap | 83.49 | 15.62 | 0.51 | 0.15 | 0.02 |
| BLEND | 58.40 | 30.53 | 6.43 | 2.62 | 1.10 |
| BLASR | 46.93 | 28.28 | 12.79 | 7.71 | 2.02 |
| bwa-mem2 | 64.27 | 22.69 | 2.62 | 1.82 | 8.18 |
| syncmer-minimap2 | 58.55 | 30.67 | 6.36 | 2.54 | 0.97 |
| minimap2 | 58.12 | 30.51 | 6.50 | 2.73 | 1.20 |
| minimap2 (old) | 58.18 | 30.51 | 6.49 | 2.71 | 1.18 |
| Magic-BLAST | 27.53 | 7.97 | 1.64 | 2.21 | 60.39 |
| Winnowmap2 | 59.59 | 30.00 | 6.08 | 2.43 | 1.02 |
| Ira | 0.93 | 10.31 | 12.69 | 29.19 | 44.77 |

**Figure 2:**
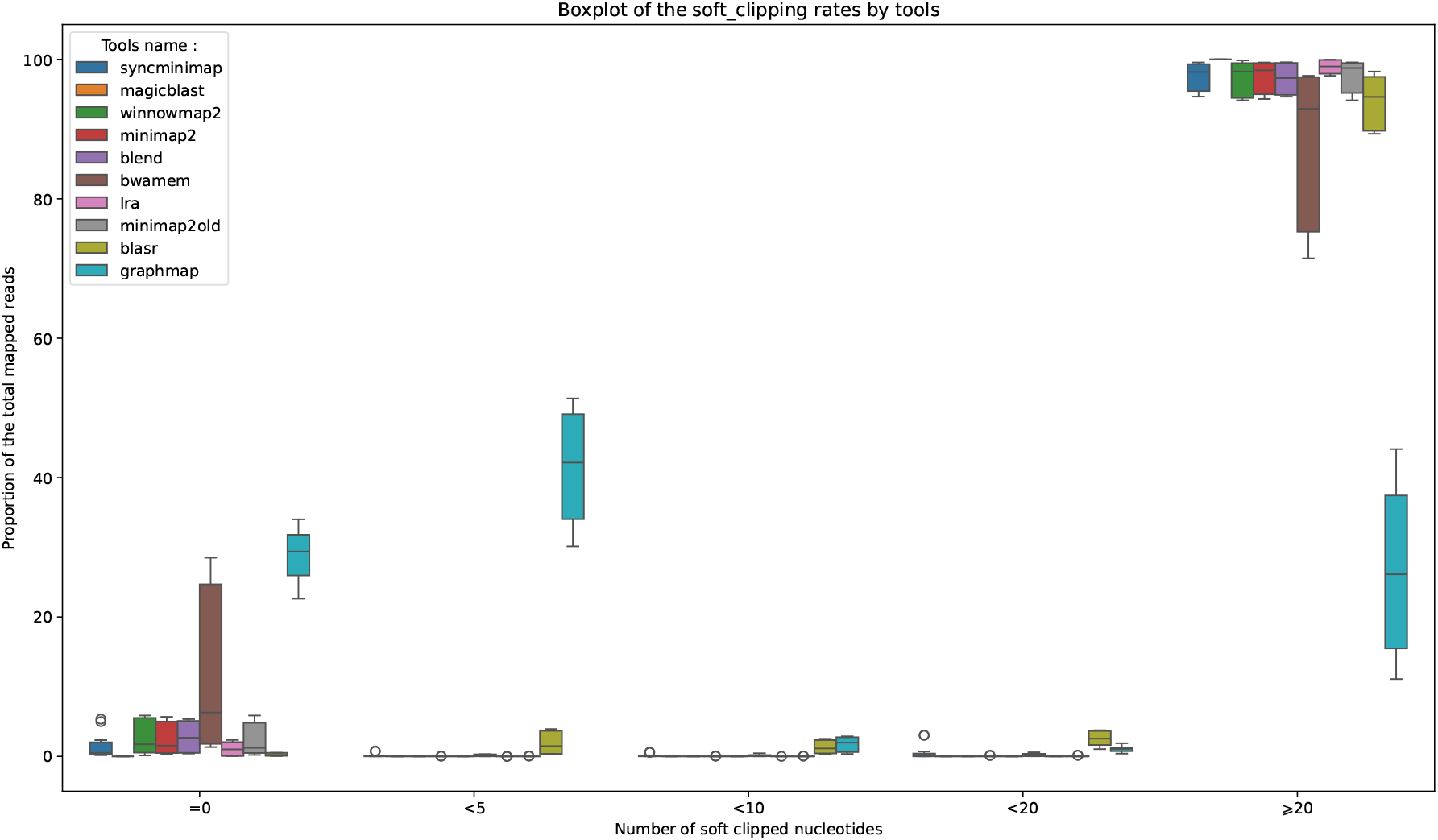
Boxplot of the percentage of soft clipping for each tool for the 34 samples generated. The x-axis show the proportion of soft clipping of the total mapped reads.

### Robustness of parametrization

Varying parameters can have more or less impact on the mapper. With mappers such as GraphMap and bwa-mem2 BLASR or Magic-BLAST, it is difficult to see any impact of the settings on the mapping. For minimap2, syncmerminimap2 and BLEND we can see a slight improvement (less than 2%) in mapper efficiency. Finally, for tools such as Winnowmap2 and lra, we can see a that some parameters improve the number of mapped reads (supplementary files S3).

### Number of indexed seeds

The number of mapped reads is positively influenced by smaller values for minimizer sizes and windows. Some tools like GraphMap index all k-mers as a regular part of their algorithm. Other, like minimap2, index a subsample of k-mers using a combination of windows and minimizers. By setting the window size and minimizer size to low values, one can be close to indexing all positions on the reference. We show for tools which natively have shorter and more numerous seeds, and for tools parametrized to harvest on almost all positions, mapping rates are better than for regular parametrization. This come at the price of an increased running time, We can see particularly for minimap2, Winnowmap2, lra, and syncmer-minimap2 that the parameter settings, which affect the number of indexed seeds, have an extremely positive impact on the number of mapped reads. For Winnowmap2, for example, we see that the median shifts from slightly below 1 to 0.

### Impact on variant calling

Each dataset includes a variant in greater or lesser proportion. In order to see if the aligned reads could correctly find the variants, we look at the different F1-scores associated with 2 different variant callers,medaka and bcftools and the F1-scores are normalized with vcfdist. It is observed in figure 3A that with medaka, despite the perfect BAM file not reaching a perfect F1-score (0.995), the vast majority of tools have an average F1-score of 1. Only GraphMap, BLASR, and Magic-BLAST have respective F1-scores of 0.914, 0.939, and 0.976. The second variant caller, bcftools, shows an F1-score around 0.5 for all tools. Figure 3B, which shows the F1-score with real data, presents completely different results. In this configuration, bcftools gives better results: 0.909 for minimap2 in both versions, 0.857 for GraphMap, 0.869 for BLASR, 0.894 for Winnowmap2, 0.871 for Magic-BLAST, 0.959 for lra, 0.902 for BLEND, 0.869 for bwa-mem2, and 0.894 for syncmer-minimap2. Variant calling with medaka is around 0.65 for most tools, except for GraphMap with 0.425, Magic-BLAST with 0.554, and bwa-mem2 with 0.554.

**Figure 3:**
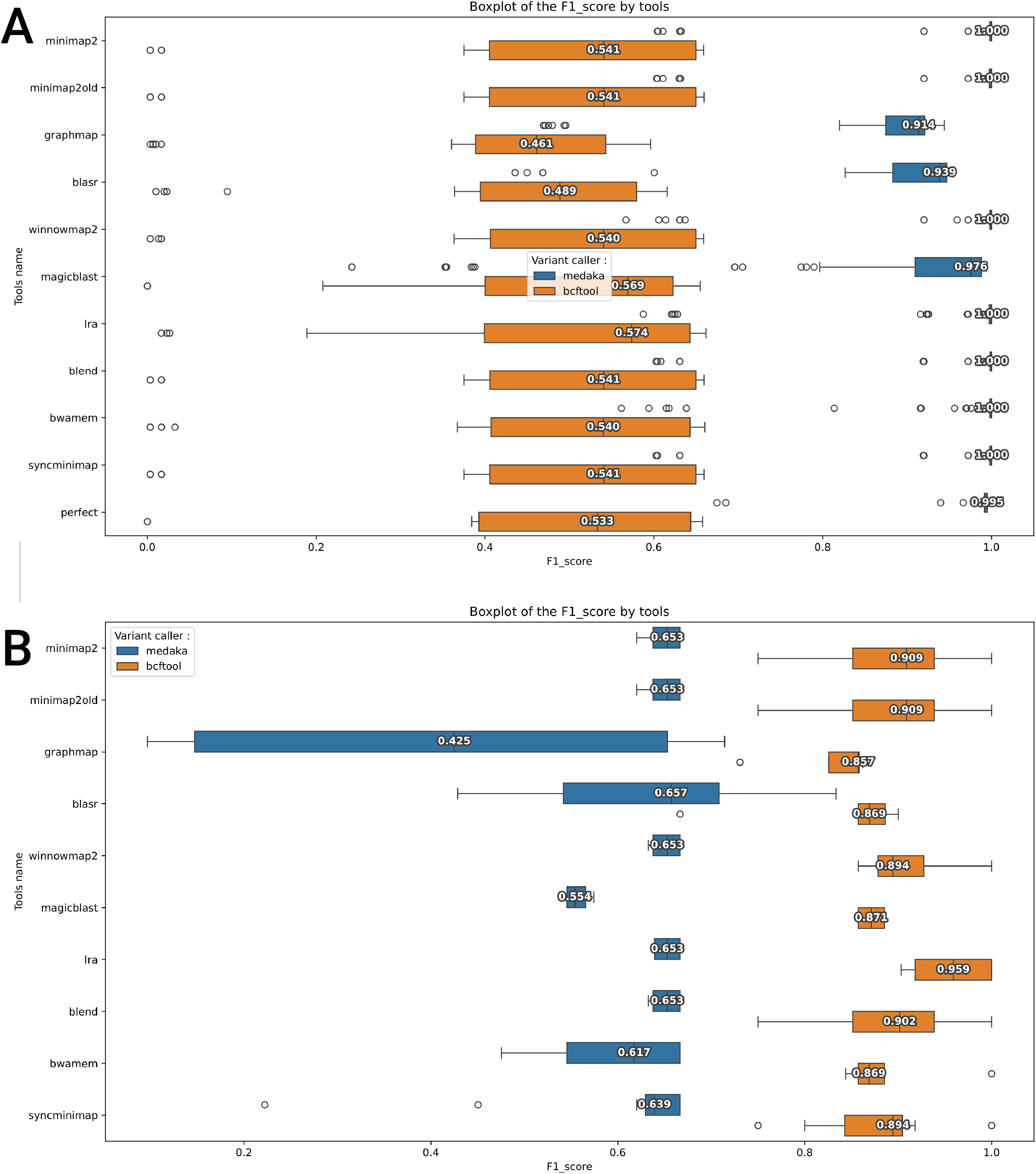
Boxplot of the F1-score for each tool across the 34 datasets. In blue, the F1-score obtained with medaka, and in orange, the F1-score obtained with bcftools.

In supplementary figures S5, which calculate the F1-score without vcfdist, different results appear with both simulated and real data. With the simulated data (figure S5A), the two variant callers have much closer F1-scores, with an average of 0.75 for bcftools and an average of 0.83 for medaka. In this configuration, Magic-BLAST with bcftools achieves the highest score, with an F1-score of 0.877.

For the real data (figure S5B), the F1-scores diverge much more between tools. With medaka, the higher F1-score is achieved by lra (0.948) and the lowest by GraphMap (0.478). For bcftools, most tools achieve an F1-score below 0.61, except for Winnowmap2 (0.675), Magic-BLAST (0.846), and lra (0.899).

#### General results regarding the different datasets

Supplementary figures S4, which show the average F1-score of all tools, indicate that there is no difference between experiments D1, D6, and D7. Therefore, it can be reasonably assumed that, for the simulated data, the choice of pore in the data generation process did not impact the results. Regarding the impact of coverage, as expected, mapping itself was not affected by it (experiments D1, D2, and D3). However, we do observe an impact on the F1-score, which will be discussed in the variant calling section. Experiment D2 suggests that, in our case, species did not influence the performance of the mappers. Although further results across more viral species would be needed to confirm this, we can still assume that viruses are less likely to show disparities in mapping results. Indeed, their small genome size leaves little room for short tandem repeat phenomena, which can be a challenge for mapping. The contamination (experiment D4) does not seem to affect the tools, which all proved to be extremely robust on the tested species, although tools like GraphMap occasionally mapped 1 or 2 human reads. Regarding read length and error rate, both factors impact the number of mapped reads. Shorter reads make mapping more difficult, likely due to lower chaining scores. This hypothesis is supported by the fact that selecting more seeds allows for a similar number of mapped reads. The error rate, as expected, affects both the number of mapped reads and the F1-score. In experiment D5, which tested mapping on a more distant species, we also observed an impact on the number of mapped reads. However, this effect is similar to what can be seen with a high error rate. Finally, the number of species present does not impact the mapping performance of the tools.

## DISCUSSION

Here, we discuss various aspects that our experiments have revealed to be important for viral read mapping.

### Soft clipping

According to the author of minimap2, clipping is the only way to report the chimeric or poor-quality reads that can be produced by long-read sequencers such as Nanopore and PacBio. In many tools, including minimap2, this implementation is not optional and it is not possible for the user to remove it. However, the implementation is not clearly indicated. minimap2’s clipping strategy can be impacted by several heuristics such as the z drop or the end bonus (we refer the reader to ^12^), but in most cases, it is during the extension part and during alignment that the clipping position will be determined by taking the positions that maximize the alignment score. This aspect of soft clipping, although it can be used to process chimeric reads or to give the mapper the possibility of managing data sets where the primers have not been removed, also represents a strong constraint on the ability to correctly retrieve information on the positioning of reads ^19^. Furthermore, no soft clipping doesn’t seem to be the best strategy either. GraphMap, which greatly minimizes the use of soft clipping, forces the alignment of barcodes attached to the reads, in the real data.

### Tool parameters

In this paper, we try to understand how the choice of a specific seed size affects the results using 3 types of parameters with a variation for the length of the reads. The various results show that parametrization has an impact on the number of reads mapped by certain tools, especially those that samples seeds. In this way, we can see that parametrizations that increase the number of seeds have a positive impact on the number of mapped reads. However, this comes at a price and will slow down calculation time. Figure S3 shows that for most tools, the default parameters are a solid choice with very good results. Despite this, for some specific samples, we can see that the use of a shorter seed improves the results. Increasing the number of seeds seems to positively affect the results independently of the types of data. The results show that decreasing the size of the seeds collected increases the results however it is important to note that decreasing the size of the reads will also increase the number of reads collected so it is more likely that the most important aspect is the number of seeds.

### Variant calling

Regarding variant calling, we observe a significant difference between the results obtained using vcf_dist, which applies corrections and filtering on the variants, and those obtained without using vcf_dist. For medaka, without vcf_dist, regardless of the origin of the BAM files, theF1-score consistently hovers around 0.8. Although the theoretically perfect BAM achieves the highest F1-score on average, the difference is only about 0.05 points compared to the average of other tools. Moreover, figure S4 shows no notable differences between the various possible parameters. However, when vcf_dist is used, we see that, apart from GraphMap, BLASR, and Magic-BLAST, all tools achieve an F1-score of 1, while the theoretically perfect BAM only scores 0.995.

In the case of bcftools, there is greater variability between mappers, and on average, magic_blast achieves the best results despite a lower mapping percentage compared to other tools, particularly for 500-bp reads. With bcftools and vcf_dist, we observe less disparity, but also a significantly lower score, with a maximum of 0.574 obtained by lra.

It is interesting to note that the F1-score for variant identification is widely used to compare mappers. While this measure provides a good indication of a mapper’s ability to process biological data, the results highlight some practical nuances. It appears that variant callers are highly specific to datasets with significant variability. Moreover, mappers with a lower average number of mapped reads like Magic-BLAST can sometimes achieve better variant calling results. Although we cannot fully explain these results at this stage, one hypothesis is that Magic-BLAST, by being very strict on the reads it maps, might only map reads with few errors, thus leading to better variant calling results compared to other mappers.

### Practices in bioinformatics

Long-read mapping tools are often designed to address specific problems at particular points in time. The first tools, such as BLASR and GraphMap, used seed types better suited to short-read data. This was not an issue when they were first developed, as long reads were relatively short. However, with the advancement of long reads, their increased length, and improved quality, new methods were introduced, most notably minimap2, which uses minimizers for mapping. One common aspect of all these tools is that they are often tested on eukaryotic and prokaryotic data. However, they all incorporate certain heuristics that, if correctly adapted to the dataset, can improve both results and implementation. Unfortunately, a general benchmark does not exist to comprehensively test and compare these tools. Although all tested tools are open source and typically provide a changelog with new releases, such a benchmark could help pinpoint improvements or degradations in their versions across different datasets.

Furthermore, as noted in this work, different tools employ various strategies for each step (seed, chain, extend), but their implementations are not modular enough to test independently and with different combinations the seeding, chaining and extending part. Finally, some post-hoc analyses become extremely specialized in using one tool in particular, such as medaka. It is very difficult to use a mapper other than minimap2 in its workflow, which has two main consequences. First, according to the way the neural network behind medaka was trained, it is likely that medaka learns from minimap2 alignments and may experience a degradation in performance with other mappers. Second, if a user wishes to use another mapper for a valid reason, it would require significant engineering effort to replace it in the workflow.

## CONCLUSION

In our experiments, we assessed whether and to which extent long-read mappers could accurately map long viral sequences. The different results obtained with the different data sets show that the tools manage to map the vast majority of reads and also manage to find the variants. However, we have sees that the particularity of viral data prevents the tools to be at their full potential with the default settings. What’s more, the great variability present in viruses can be a real challenge if the species sequenced are too far from the reference species. We have shown that recent or well-maintained tools, such as minimap2 or lra, are generally preferable for aligning viral reads. When precision is needed at the edges of the alignment - as in the search for subgenomic RNAs ^19^ or splicing events ^34^ - BWA-mem can serve as a valuable middle ground, minimizing the rate of soft clippings without excess.

We have also demonstrated that an interaction link exists between mappers and downstream tools. Thus, when a mapper appears to align a greater quantity of reads effectively, the details of the alignment can either favor or hinder variant calling tools, depending on how they have been trained.

## AVAILABILITY

The benckmark used to generate the results can be found at https://github.com/ThomasBaudeau/BenchMapL

## ACKNOWLEDGMENTS

This work was funded by ANR INSSANE ANR21-CE45-0034 project.

## AUTHOR CONTRIBUTIONS

TB, MS and CM designed the experiments and wrote the paper. TB conducted the experiments.

