## Supplementary for "Assessing Long-Read Mappers for Viral Genomics"

### 1 Additional Figures

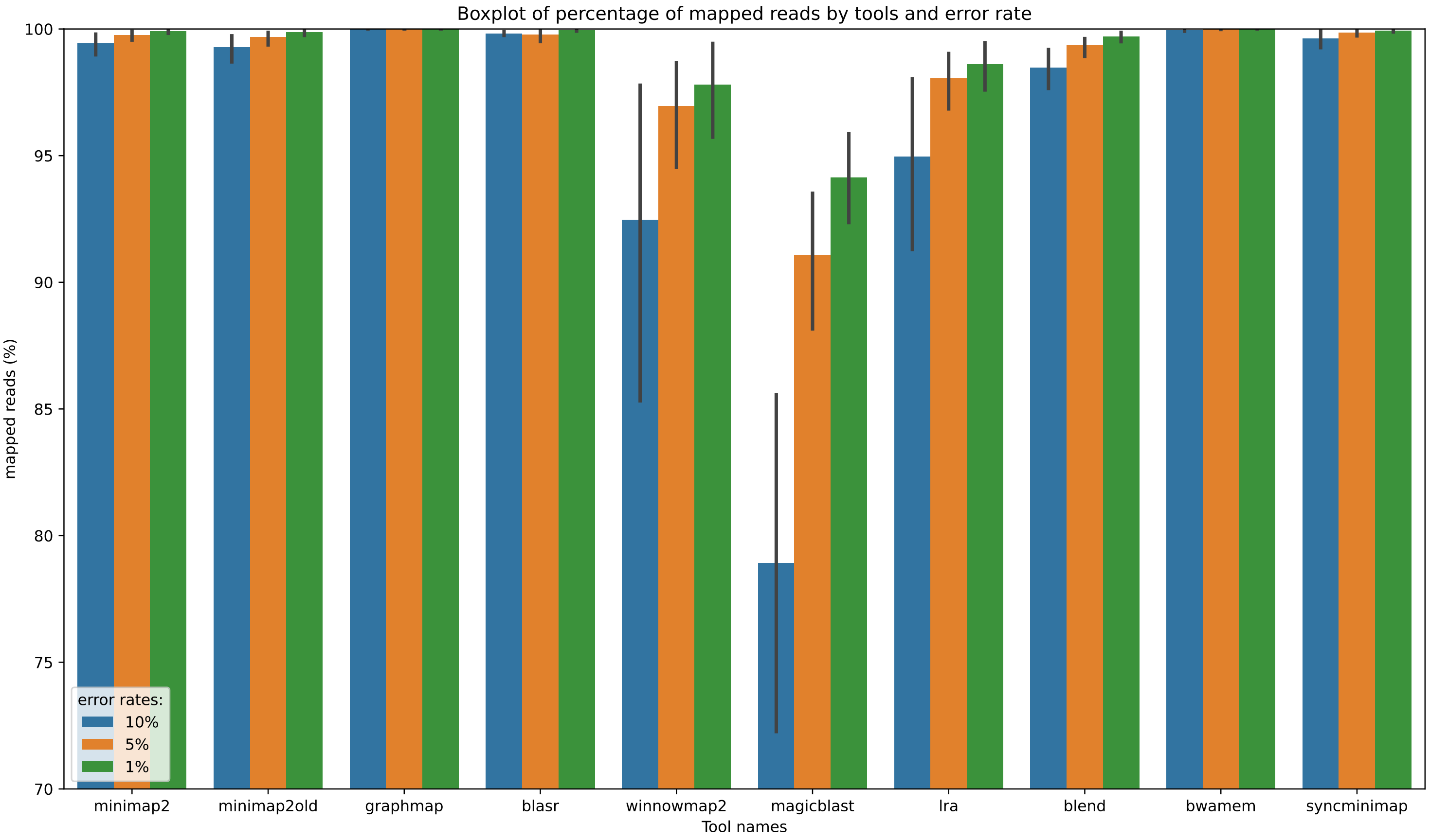


Figure S1: Boxplot of the percentage of mapped reads by tools and reads error rates

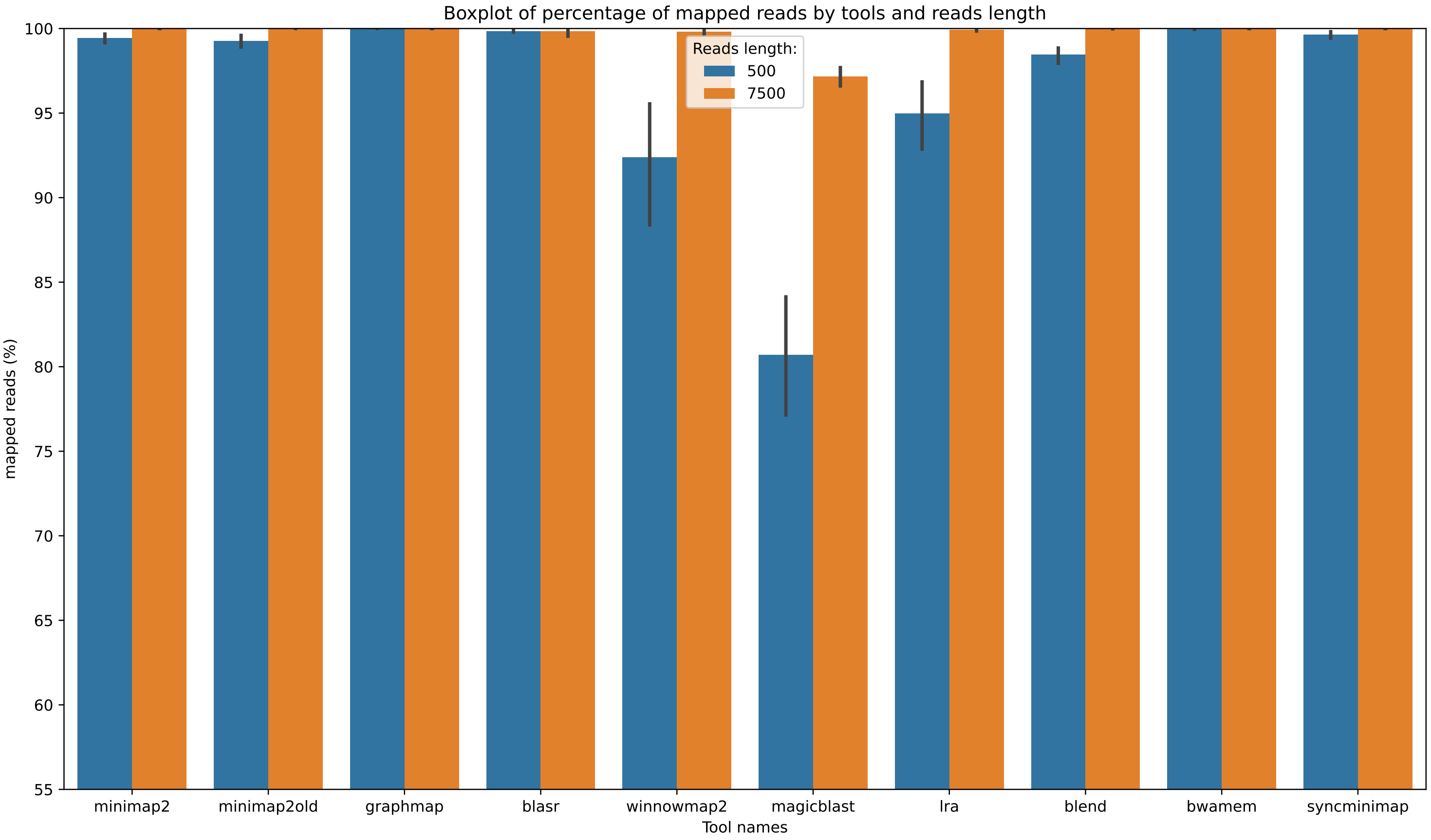


Figure S2: Boxplot of the percentage of mapped reads by tools and reads length

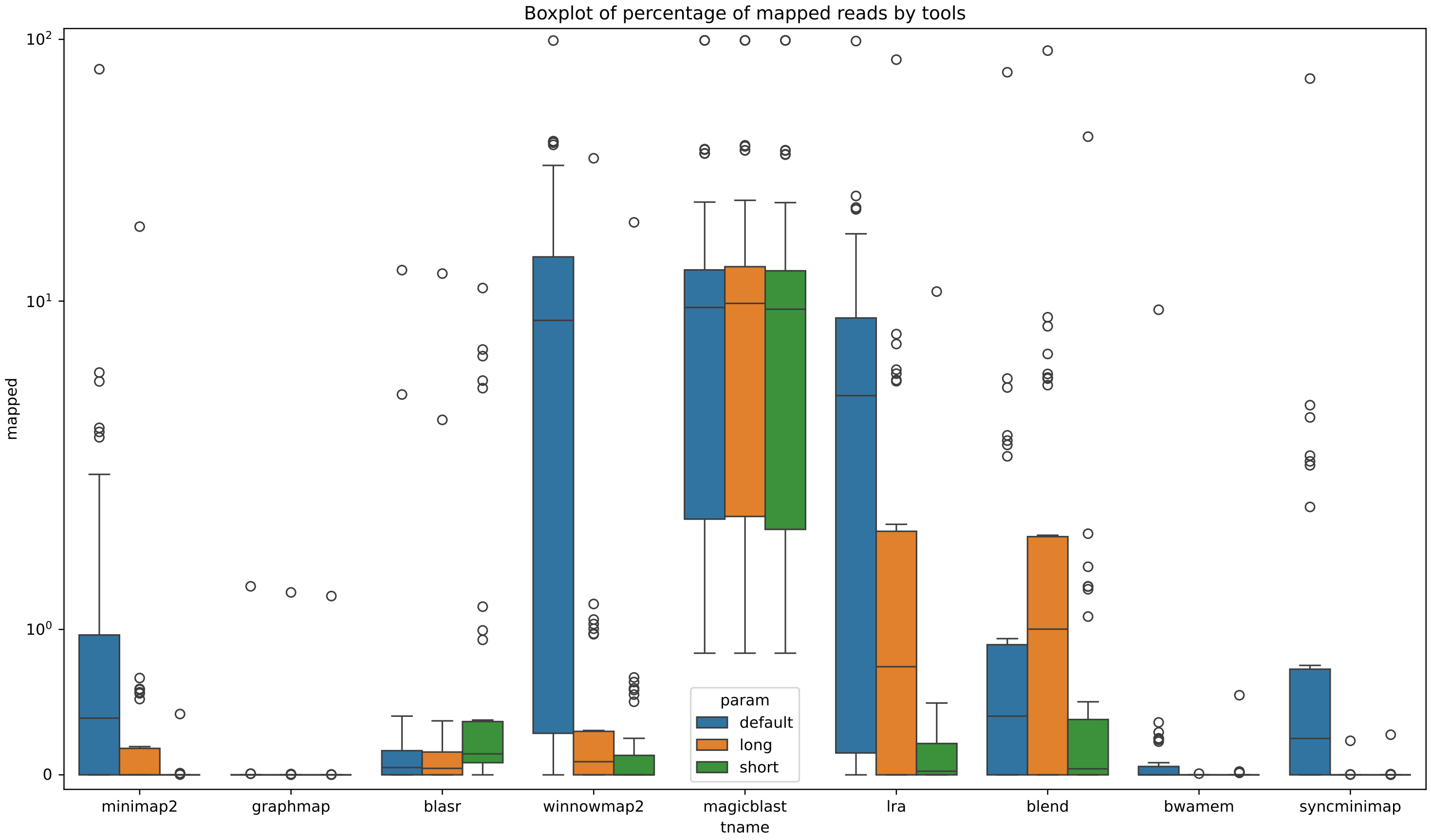


Figure S3: Boxplot of the % of unmapped reads by tools and parameters

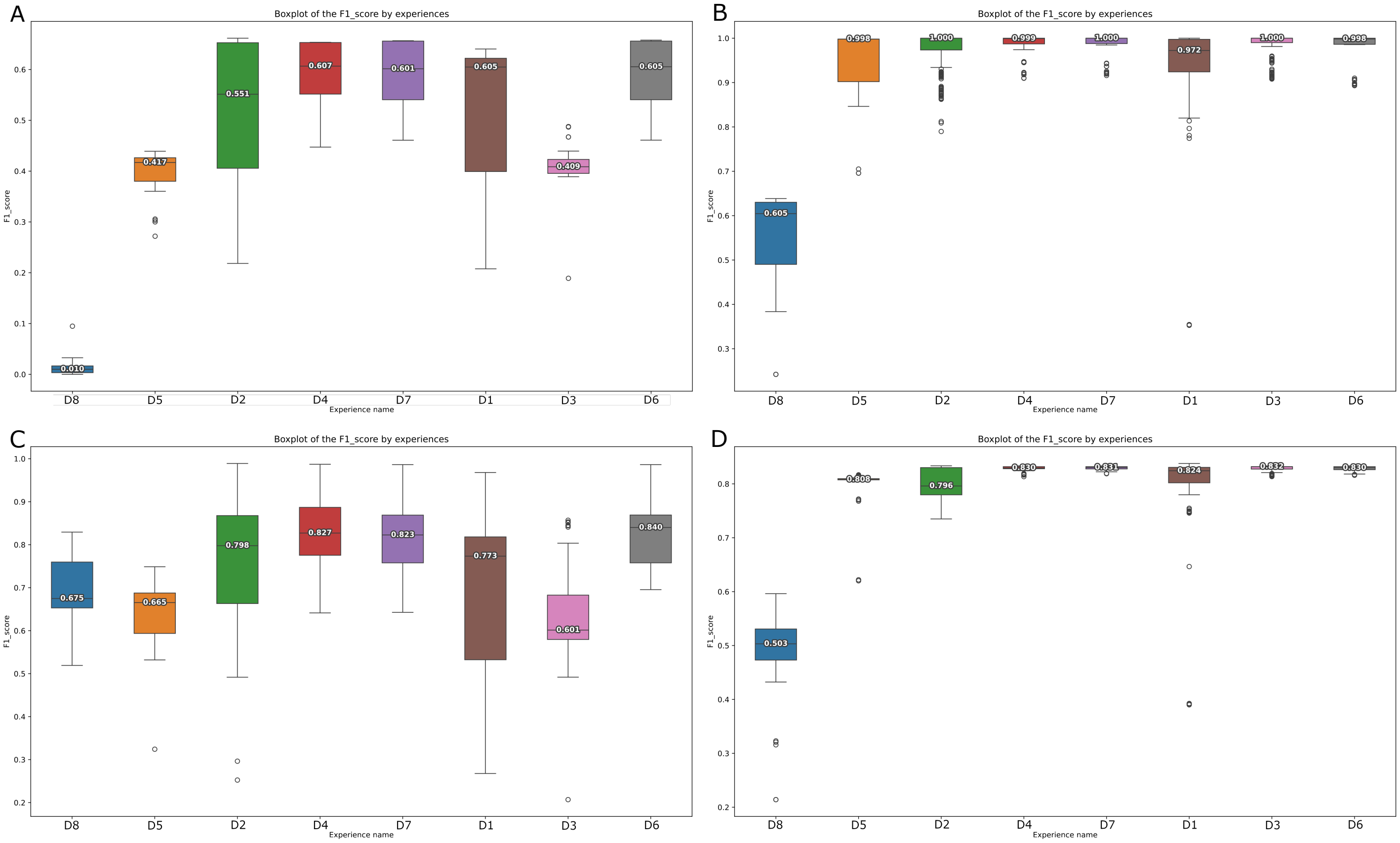


Figure S4: Boxplot of the F1-scores by experiences, A) bcftools F1-scores with vcfdist; B) medaka F1-scores with vcfdist; C) bcftools F1-scores without vcfdist; D) medaka F1-scores without vcfdist;

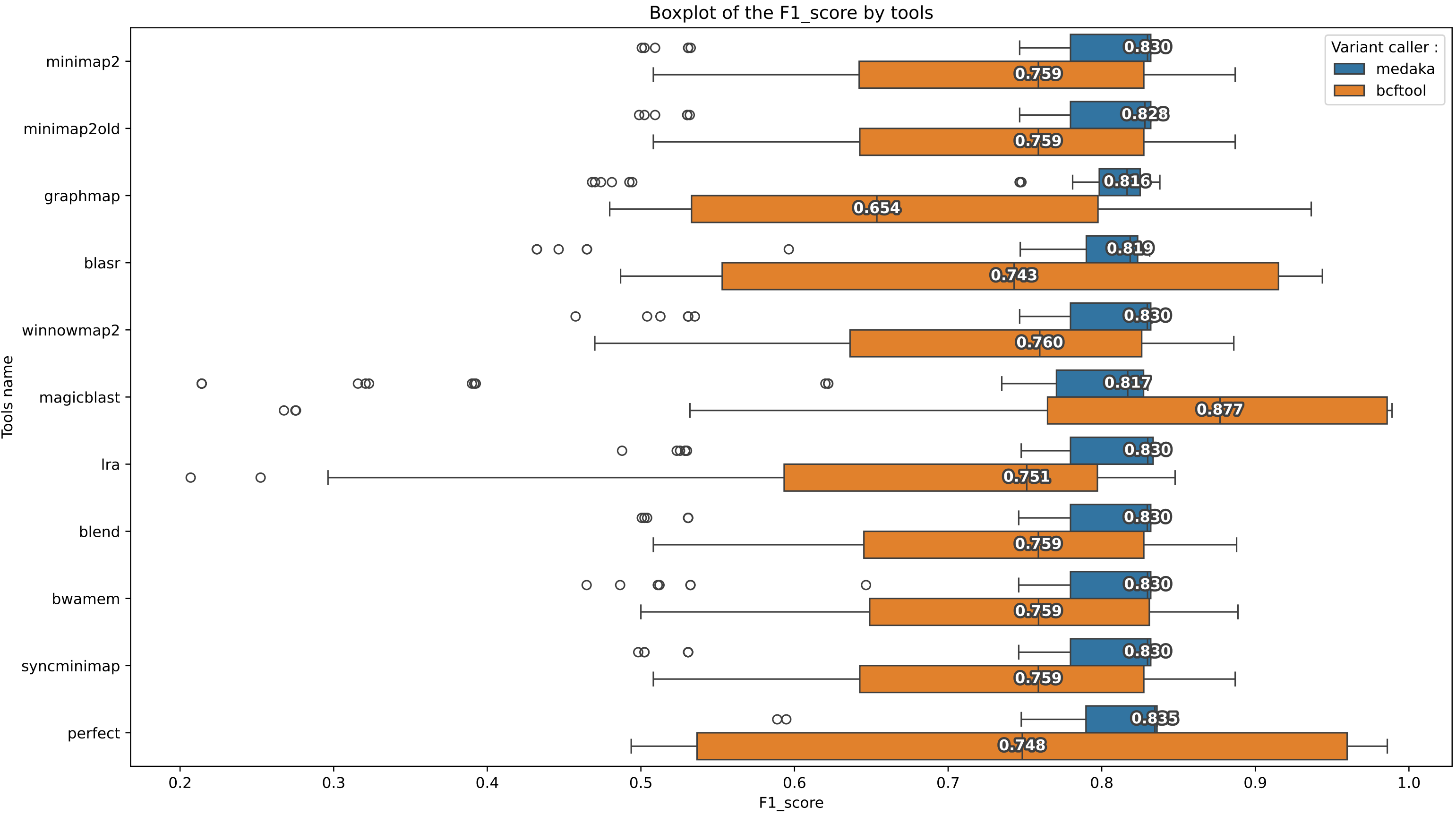


Figure S5: Boxplot of the F1-score by tools without using vcfdist

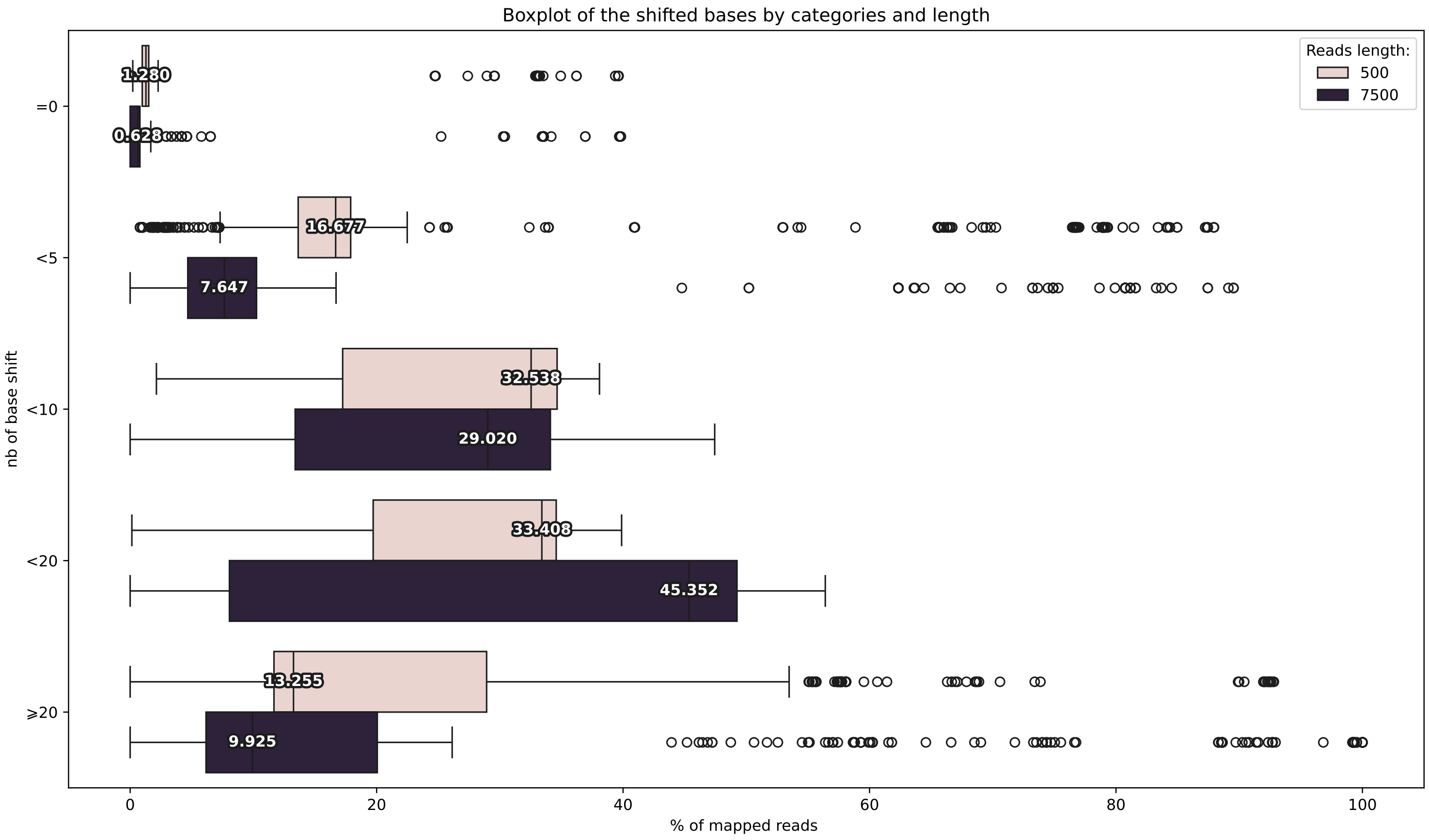


Figure S6: Boxplot of the percentage of shifted bases per categories and error rates

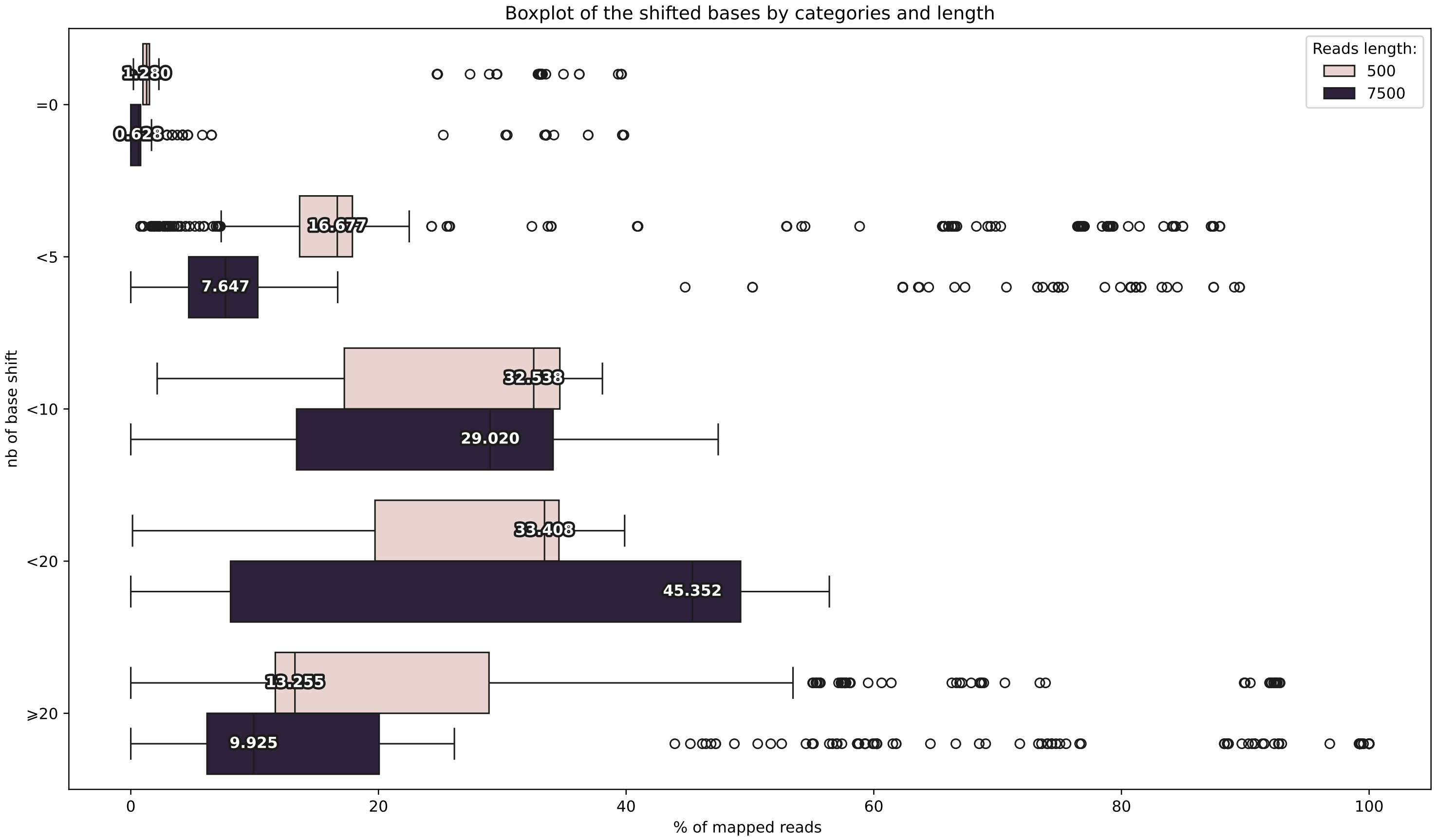


Figure S7: Boxplot of the percentage of shifted bases per categories and length
